# Tryptophan specialized metabolism and ER body-resident myrosinases modulate root microbiota assembly

**DOI:** 10.1101/2022.07.06.498822

**Authors:** Arpan Kumar Basak, Anna Piasecka, Jana Hucklenbroich, Gözde Merve Türksoy, Rui Guan, Pengfan Zhang, Felix Getzke, Ruben Garrido-Oter, Stephane Hacquard, Kazimierz Strzałka, Paweł Bednarek, Kenji Yamada, Ryohei Thomas Nakano

**Author notes:** To whom correspondence may be addressed: Ryohei Thomas Nakano, Max Planck Institute for Plant Breeding Research. The Medical Research Council Toxicology Unit, University of Cambridge, Cambridge, UK.

## Abstract

Indole glucosinolates (IGs) are tryptophan (Trp)-derived sulfur-containing specialized metabolites that play a crucial role in plant-microbe interactions in plants of the order Brassicales, including *Arabidopsis thaliana*. Despite the growing body of evidence implicating IG biosynthetic pathways in root-microbiota interactions, how myrosinases, the enzymes that convert inert IGs into bioactive intermediate/terminal products, contribute to this process remains unknown. Here, we describe the roles of the PYK10 and BGLU21 myrosinases in root-microbiota assembly partly via metabolites secreted from roots into the rhizosphere. PYK10 and BGLU21 localize to the endoplasmic reticulum (ER) body, an ER-derived organelle observed in plants of the family Brassicaceae. We investigated the root microbiota structure of mutants defective in the Trp metabolic (*cyp79b2b3* and *myb34/51/122*) and ER body (*nai1* and *pyk10bglu21*) pathways and found that these factors together contribute to the assembly of root microbiota. Microbial community composition in soils as well as in bacterial synthetic communities (SynComs) treated with root exudates axenically collected from *pyk10bglu21* and *cyp79b2b3* differed significantly from those treated with exudates derived from wild-type plants, pointing to a direct role of root-exuded compounds. We also show that growth of the *pyk10bglu21* and *cyp79b2b3* mutants was severely inhibited by fungal endophytes isolated from healthy *A. thaliana* plants. Overall, our findings demonstrate that root ER body-resident myrosinases influencing the secretion of Trp-derived specialized metabolites represent a lineage-specific innovation that evolved in Brassicaceae to regulate root microbiota structure.

**Significance:** ER bodies were first identified in roots of Brassicaceae plants more than 50 years ago, but their physiological functions have remained uncharacterized. A series of previous studies have suggested their possible role in root-microbe interactions. Here, we provide clear experimental evidence showing a role for ER bodies in root-microbiota interactions, which overlaps with that of root-exuded Trp-derived metabolites. Our findings delineate a plant lineage-specific innovation involving intracellular compartments and metabolic enzymes that evolved to regulate plant-microbe interactions at the root-soil interface.

## INTRODUCTION

ER bodies (Fig 1A; 1) are spindle-shaped organelles that have evolved in plants from taxonomically related families in the order Brassicales, i.e., Brassicaceae, Cleomaceae, and Capparaceae (2, 3). These compartments were originally named “dilated cisternae” upon their discovery using an electron microscope (4), and, since then, a half-century of studies has revealed the molecular nature of these structures. ER bodies in *Arabidopsis thaliana* accumulate large concentrations of β-glucosidase enzymes, PYK10 and BGLU21 in roots/seedlings and BGLU18 in rosette leaves (5–7). A basic helix-loop-helix (bHLH) transcription factor NAI1 controls the formation of ER bodies by positively regulating the expression of genes encoding ER body constituents, including PYK10 and BGLU21, a scaffold protein NAI2, and integral membrane proteins MEMBRANE OF ER BODY 1 (MEB1) and MEB2 (5, 8–13).

**Figure 1.**
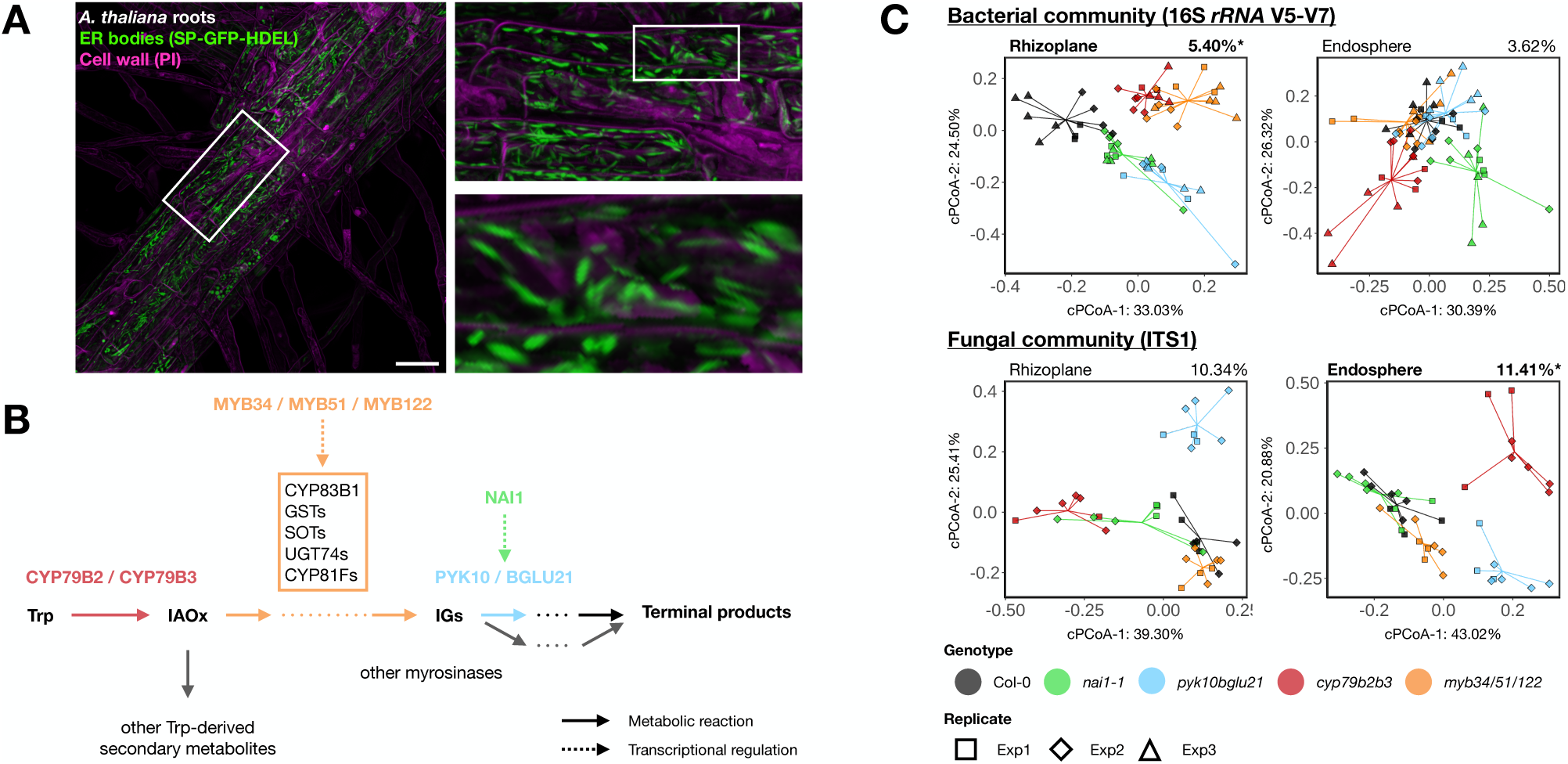
ER bodies and Trp metabolism play a role in root-associated microbiota assembly. (*A*) Roots of *A. thaliana* developing a large amount of ER bodies, visualized by ER-localized GFP (SP-GFP-HDEL). Cell wall was stained by propidium iodide. The bar corresponds to 50 μm. (*B*) Representation of IG biosynthetic and catabolic pathways. Arrows and dotted arrows indicate metabolic reactions and transcriptional regulation, respectively. (*C*) Constrained principal coordinates analysis (PCoA) of the bacterial and fungal community structures in the roots of Col-0 as well as mutants impaired in ER body formation (*nai1-1*), ER body-accumulating myrosinases (*pyk19bglu21*), IG biosynthesis (*myb34/51/122*), and Trp metabolism (*cyp79b2b3*) based on Bray-Curtis dissimilarities. Ordination was constrained by genotypes and conditions by soil batches, biological replicates and sequencing runs. Colours and shapes represent the genotypes and biological replicates, respectively. Variation explained by genotypes based on permutational analyses of variance (PERMANOVA; n = 999) is indicated at top-right. Asterisks indicate statistical significance based on PERMANOVA (*α* = 0.05). Trp, tryptophan; IAOx, indole-3-acetoaldoxime; IG, indole glucosinolates; GST, glutathione-*S*-transferase; SOT, sulfotransferase; UGT, uridine diphosphate-glycosyltransferase.

ER bodies are constitutively developed in roots and seedlings, while they are strongly induced, both locally and systemically, by wounding or jasmonic acid treatment in mature rosette leaves (7, 14). Recent studies have described the role of leaf ER bodies in defence against herbivores (woodlice), likely mediated by the β-glucosidase (BGLU) activity of PYK10 and BGLU18 (6, 15). In particular, these BGLUs mediate the hydrolysis of glucosinolates (6, 16), a class of specialized metabolites (also known as secondary metabolites) that accumulate in plants of the order Brassicales and that are crucial for defence against herbivores and microbial pathogens (17). The enzymatic capacity to hydrolyse the thioglucosidic bond in glucosinolates is called myrosinase activity and is harboured by only a subset of BGLUs (18). Myrosinase activity toward the tryptophan (Trp)-derived group of glucosinolates with an indole side chain (indole glucosinolates, IG) has been demonstrated for PYK10 and BGLU18 (6, 16), while, *in vitro*, their specific activity towards sinigrin, a short-chain methionine (Met)-derived aliphatic glucosinolate (AG), appears to be lower (5, 19). On the other hand, PYK10 and BGLU21 can hydrolyse AGs in plant tissue homogenates (15), suggesting that these ER body myrosinases can act on a wide range of glucosinolates. Overall, these studies suggested a role of ER Bodies related to the metabolism of glucosinolates.

In addition to their well-characterized role in restricting pathogenic infection in leaves (20–23), root-accumulating Trp-derived metabolites, including IGs, play a crucial role in interaction with soil-borne non-pathogenic microbes at the root-soil interface (24–28). For example, the *cyp79b2b3* double mutant, in which the major branch of Trp metabolism (the metabolic pathway initiated with a conversion of Trp into indole-3-acetoaldoxime, IAOx) is entirely blocked, is unable to properly manage the accommodation of beneficial endophytic fungi and exhibits severe growth inhibition in the presence of these microbes (25, 27). The *cyp79b2b3* mutant also fails to control the overall abundance of fungi in roots (“fungal load”), resulting in a severe growth defect when inoculated with a synthetic community (SynCom) composed of commensal bacteria and endophytic fungi (28). Notably, Trp-derived specialized metabolites including IG catabolites are reported to be secreted into the rhizosphere (29–31), and an alteration in the profiles of root-produced glucosinolates has an impact on the root-associated bacterial and fungal microbiota (32); whether intact IGs are also secreted along with the IG catabolites appears to be dependent on the investigated species and/or the experimental conditions. Based on these findings, it has been proposed that root ER bodies are involved in plant defence and/or interaction with surrounding microbes via Trp metabolism (16, 33). However, to date, the specific role for the root ER bodies under natural conditions remains unclear.

We hypothesized that the PYK10-mediated hydrolysis of Trp-derived specialized metabolites, including IGs, produces a variety of terminal products and plays an important role in plant-microbiota interactions at the root-soil interface. In this study, we provide evidence that ER bodies and Trp-derived metabolic pathways influence root microbiota composition in an overlapping manner. We also show that this effect on microbial communities is partially mediated by root-exuded compounds and has a direct impact on bacterial community composition in the absence of fungi, as well as on fungal behaviour in the absence of bacteria. Overall, our results demonstrate a physiological role for root ER bodies in shaping root microbiota assembly at the root-soil interface.

## MATERIALS AND METHODS

### Plant, microbial, and soil materials

Cologne agricultural soil batches 11 (CAS11), 13 (CAS13) and 15 (CAS15) were harvested in February 2017, February 2019, and in January 2020, respectively, as described previously (34). *Arabidopsis thaliana* wild-type Col-0 seeds were obtained from the Nottingham Arabidopsis Stock Centre (NASC). The *pyk10-1 bglu21-1* double mutant (*pyk10bglu21*), *nai1-1*, *myb34 myb51 myb122* triple mutant (*myb34/51/122*) and *cyp79b2 cyp79b3* double mutant (*cyp79b2b3*) plants have been described previously (5, 35–37). The bacterial and fungal strains used in this study (Table S1 and S2) were described previously (25, 38, 39).

### Root harvesting and fractionation

After cultivation of plants, roots and soils were harvested from pots and fractionated into rhizosphere, rhizoplane, and endosphere fractions, as described previously (40). Briefly, the soil particles physically attached to the root surface were collected as a rhizosphere fraction by shaking vigorously in sterile water followed by centrifugation. Microbes on the root surface were collected as a rhizoplane fraction by washing roots with detergent followed by filtration through a 0.22-µm membrane. Roots were then surface-sterilized with 70% (v/v) ethanol and bleach solution to give the endosphere fraction. All samples were immediately frozen in liquid nitrogen and stored at −80 °C until processing.

### Collection of root exudates

Root exudates were collected from plants hydroponically grown in axenic glass jars containing glass beads, as described previously (41). Seeds of Col-0 wild-type as well as *pyk10bglu21* and *cyp79b2b3* mutant plants were surface-sterilized with 70% (v/v) ethanol and bleach with 12% (w/v) active chlorine and germinated on metal meshes placed on half-strength Murashige and Skoog (MS) basal salts (Sigma-Aldrich), supplemented with 1% (w/v) sucrose and 1% (w/v) agar. Seeds were stratified for 48 hours at 4 °C in the dark and cultured for four days under short-day conditions (10 hours under light at 21 °C and 14 hours under dark at 19 °C). The four-day-old seedlings on the mesh were transferred aseptically into glass jars containing sterile glass beads (1 mm) and 26 ml of half-strength liquid MS media. Glass jars were placed in a breathable Microbox (Sac O2, Belgium) and cultivated for five weeks under short-day conditions. Hydroponic medium was collected using an aseptic stainless needle into 50-mL tubes and concentrated to 1/10 volume by a lyophilizer.

### Metabolite profiling of root exudates

The LC-MS system consisted of UPLC with a photodiode-array detector PDAeλ (Acquity System; Waters) hyphenated with a high-resolution QExactive hybrid MS/MS quadrupole Orbitrap mass spectrometer (Thermo Scientific, http://www.thermofisher.com/). Chromatographic profiles of metabolites and the quantitative measurements were obtained using water acidified with 0.1% formic acid (solvent A) and acetonitrile (solvent B) with a mobile phase flow of 0.35 ml min^-1^ on an ACQUITY UPLC HSS T3 C18 column (2.1 × 50 mm, 1.8 μm particle size; Waters) at 22 °C. The root-exudate sample (5 μL) was injected into the inlet port after purging and rinsing the system. The active MS was operated in Xcalibur version 3.0.63 with the following settings: heated electrospray ionization ion source voltage −3 kV or 3 kV; sheath gas flow 30 L min^-1^; auxiliary gas flow 13 L min^-1^; ion source capillary temperature 250 °C; auxiliary gas heater temperature 380 °C. MS/MS mode (data-dependent acquisition) was recorded in negative and positive ionization, in resolution 70000 and AGC (ion population) target 3e+6, scan range 80 to 1000 mz. Obtained LC-MS data were processed for peak detection, deisotoping, alignment and gap-filling by MZmine 2.51 (42) separately for positive and negative ionization mode, then data from both modes were combined. The prepared data table was post-processed for missing values imputation, log transformation and data filtering by MetaboAnalyst (43). Then, data were visualized by Sparse Partial Least Squares Discriminant Analysis and Venn diagrams showing signals selected by one-way ANOVA with Benjamini-Hochberg correction with the following criteria: FDR ≤ 0.05, and | log fold change | ≥ 1.0), where fold change is calculated for specific comparison of *cyp79b2b3* vs Col-0 and *pyk10bglu21* vs Col-0. Glucosinolates were identified based on the MS/MS fragmentation spectra. Standard compounds were used for identification and quantification of IAA, esculetin, fraxetin and scopoletin. Pathway enrichment analysis of the root exudate metabolome was conducted by the MetaboAnalyst modules “Functional Analysis” followed by “Pathway enrichment” on signals selected in previous ANOVA and fold change analysis. Annotation of signals to the KEGG pathway for *A. thaliana* was conducted with a mass tolerance of 5 ppm.

### Collection of root extracts

To collect root extracts, we cultured Col-0 wild-type, as well as *pyk10bglu21* and *cyp79b2b3* mutant plants on half-strength MS media, supplemented with 1% (w/v) sucrose and 1% (w/v) agar for 21 days. Roots were weighed, harvested and frozen in liquid nitrogen and stored at - 80 °C until further processing. Immediately before treatment, roots were homogenized in phospho-buffered saline using 1-mm zirconia beads to obtain 10 mg/mL of root extracts.

### Soil treatment with root exudates

Approximately 500 mg of CAS soil were transferred into 2 mL screw-cap tubes and treated with 50 µL of root exudates or root extracts. Tubes were covered by breathable tape and incubated at 28 °C under dark conditions. Root exudates or freshly prepared root extracts were added every three days to replenish the weight lost due to evaporation of the moisture content. Soils were freeze-dried and stored at −80 °C until further processing.

### Bacterial microbiota reconstitution experiment

Bacterial cultures were retrieved from glycerol stocks on plates containing 50% (w/v) trypsin soy broth (TSB) supplemented with 1.5% (w/v) agar. Single colonies were used to inoculate 50% TSB (15 g/L) liquid media in 96-deep-well plates and cultured for four days at 25 °C. Sixty microlitres of cultures were transferred to another 96-well plate containing 600 µL of the sterile TSB media cultured for another three days in parallel to the original culture plates. Cultures from 7-day-old and 3-day-old plates were pooled and OD_600_ was adjusted to roughly 0.5 for each strain. Individual strains were pooled into a new tube to give a final bacterial concentration of OD_600_ = 1. The bacterial inoculum was centrifuged at 3,000 rpm for 10 minutes to remove the TSB, followed by two washes in 10 mM MgCl_2._ The obtained pellet was dissolved in MgCl_2_ and OD_600_ was adjusted to 1. The starting inoculum was incubated overnight at 25 °C, such that bacteria would have consumed nutrients carried over from previous cultures and utilize the root exudate as their sole nutrient source during the treatment. On the day of the SynCom treatment, the concentration of the starting inoculum was adjusted to OD_600_ 0.5. The inoculum was diluted to OD_600_ 0.05 and 0.005 in 50 µl of root exudates. We multiplexed 1,216 samples in two MiSeq runs with barcode sequences in both forward and reverse primers (44) and used a reference-based error correction algorithm (Rbec; 45) to maximize the number of reads used for the quantification of their abundance. At the time of harvest (24 or 72 hours post inoculation; 24 or 72 hpi), a fixed amount of *Escherichia coli* DH5*α* cells (at final OD_600_ of 0.01 or 0.001 depending on the initial inoculum titre), whose 16S *rRNA* is fully distinguishable from all strains included in our SynCom, was added to each sample to enable quantitative abundance (QA) estimation (46). The harvested cultures were immediately processed for DNA isolation.

### DNA extraction and amplicon sequencing

Total DNA from the rhizosphere, rhizoplane and root samples was extracted using the FastDNA SPIN Kit for Soil (MP Biomedicals, Solon, USA), as described previously (40). Bacterial cells in liquid cultures were lysed in alkaline sodium hydroxide and directly used as a PCR template, as described previously (38). The V5–V7 region of bacterial *16S* rRNA gene and the ITS1 region of fungal DNA was PCR-amplified by specific primer sets containing adapter sequences for sequencing and barcode sequences for multiplexing (Tables S3–6). Approximately the same amounts of PCR product were pooled, purified twice using AMPure XP beads (Agencourt) and sequenced by an Illumina MiSeq platform (MiSeq Reagent Kit V3, 600-cycle).

### Bioinformatic analysis of microbiome profiling

Pre-processing, demultiplexing and analysis of amplicon sequence variants (ASVs) was performed as described previously (47) using the DADA2 pipeline (48). Taxonomic assignments for ASVs were performed referring to the SILVA (v138) database for bacteria and UNITE (release 04.02.2020; 49) database for fungi. Reference-based analysis of bacterial SynCom data was performed using the Rbec pipeline (45).

### Statistical analysis for microbiome profiling

All statistical analyses were performed in R (https://www.r-project.org/). Unconstrained and constrained principal coordinates analyses (PCoA and CPCoA) were performed based on Bray-Curtis dissimilarities using the cmdscale and capscale functions in the *stats* and *vegan* packages. Differential abundance of ASVs and aggregated ASVs were conducted using the *edgeR* package by fitting relative abundance to a generalized linear model with a negative binomial distribution, controlling for sequencing batch, technical replicates and the experimental batch as random factors.

### Plant-fungi binary interaction assay

Fungal inoculation was performed as described previously (24, 39). About 50 mg of fungal mycelium were collected and homogenized in 10 mM MgCl_2_, which was then used to inoculate surface-sterilized seeds. Plants and fungi were co-cultured on half-strength MS media supplemented with 1% (w/v) agar for 21 days under short-day conditions, and their shoot fresh weights were measured after cultivation.

## RESULTS

### ER bodies and Trp-derived specialized metabolites together contribute to root microbiota assembly

To investigate the impact of ER bodies and Trp-derived specialized metabolites on root microbiota assembly, we performed amplicon sequencing analysis of bacterial (16S *rRNA*) and fungal (ITS1) microbiota community compositions of *A. thaliana* mutant roots (rhizoplane and endosphere fractions) impaired in ER body-resident myrosinases (*pyk10bglu21*), the formation of ER bodies (*nai1-1*), the biosynthesis of IGs as well as other Trp-derived specialized metabolites to a milder extent (*myb34/51/122*), and the entire specialized Trp metabolism (*cyp79b2b3*), along with the Col-0 wild-type plants (Fig. 1B), grown in natural soils (34). The analysis of *β*-diversity at the amplicon sequence variant (ASV) level revealed significant differences between the plant genotypes in the bacterial communities in the rhizoplane (*P* = 0.009; 5.40% of variation explained by genotypes) and in the fungal communities in the endosphere (*P* = 0.021; 11.41%), while bacterial and fungal communities in the endosphere and rhizoplane, respectively, did not significantly differ across genotypes (Fig 1C; Table S7). Bacterial communities in the rhizoplanes of *cyp79b2b3* and *myb34/51/122* and of *pyk10bglu21* and *nai1* were similar to each other, respectively (Fig 1C; top left), pointing to an active role of IGs and ER bodies in bacterial community assembly rather than a stochastic variation between different host genotypes. On the other hand, in fungal communities, only *cyp79b2b3* and *pyk10bglu21* mutants, in which enzyme-encoding genes were disrupted, showed substantial differences from the wild type, while *myb34/51/122* and *nai1*, the transcription factor mutants, exhibited a milder impact (Fig. 1C; bottom right). This may be explained by Trp-derived metabolites other than IGs accumulating to levels similar to Col-0 and/or residual amounts of IGs and PYK10 and BGLU21 in these mutant roots (9, 22). A similar trend of microbial community shifts in mutants compared to the wild type was observed when we aggregated the relative abundance (RA) of ASVs at the family level (Fig S1); this analysis showed that the observed community shift occurred at the family or higher taxonomic level, rather than at the strain- or species-specific level.

We noted that the bacterial community shift observed in the rhizoplane of mutants compared to the wild type was in the same direction along the first axis, while the ER body mutants and Trp metabolism mutants were separated along the second axis (Fig. 1C; top left). This implied that the loss of ER bodies and Trp-derived metabolites exerted a partly similar impact, if not entirely identical, on the root-associated microbiota structure. This was also supported by the significant positive correlation between the log_2_-scale fold changes (logFC) in RA values of ASVs or families between each mutant and wild-type plants (Fig S2 and S3). Namely, bacteria belonging to the families Burkholderiaceae and Oxalobacteraceae/Xanthomonadaceae appeared to be commonly enriched and depleted, respectively, in the mutant rhizoplane compared to the wild type (Fig. S4A). Likewise, fungi belonging to the families Nectriaceae, Plectosphaerellaceae and Cladosporiaceae were found to be commonly enriched in the mutant endosphere compared to the wild type (Fig S4B). Overall, these data demonstrate that the ER body-resident myrosinases and Trp metabolism contribute to microbial community assembly, affecting the behaviour of overlapping sets of microbes.

### Trp-derived metabolites secreted into the rhizosphere in a manner dependent on PYK10 and BGLU21 modulate microbiota community structure

The significant bacterial community shift at the rhizoplane (root exterior) suggested a role for root-exuded compounds in bacterial community assembly. To directly test this, we collected root exudates from Col-0 wild-type as well as *pyk10bglu21* and *cyp79b2b3* mutant plants, axenically grown in a hydroponic culture system (41). We first analysed the collected exudates by untargeted HPLC-MS/MS, which revealed a significant difference in the compositions of metabolites between exudates collected from mutants and the wild type (Fig. 2A and Fig S5; Table S8). We observed up to 696 and 1,307 signals with significant differences in their RA values in *pyk10bglu21* and/or *cyp79b2b3* root exudates compared to Col-0 exudates, among which 34 and 273 signals were commonly depleted/enriched in both mutant exudates (Fig. 2B and Table S9). This result demonstrates that Trp-derived specialized metabolites are indeed secreted to the rhizosphere, and PYK10 and BGLU21 either directly or indirectly contribute to the secretion of these metabolites as well as other metabolites.

**Figure 2.**
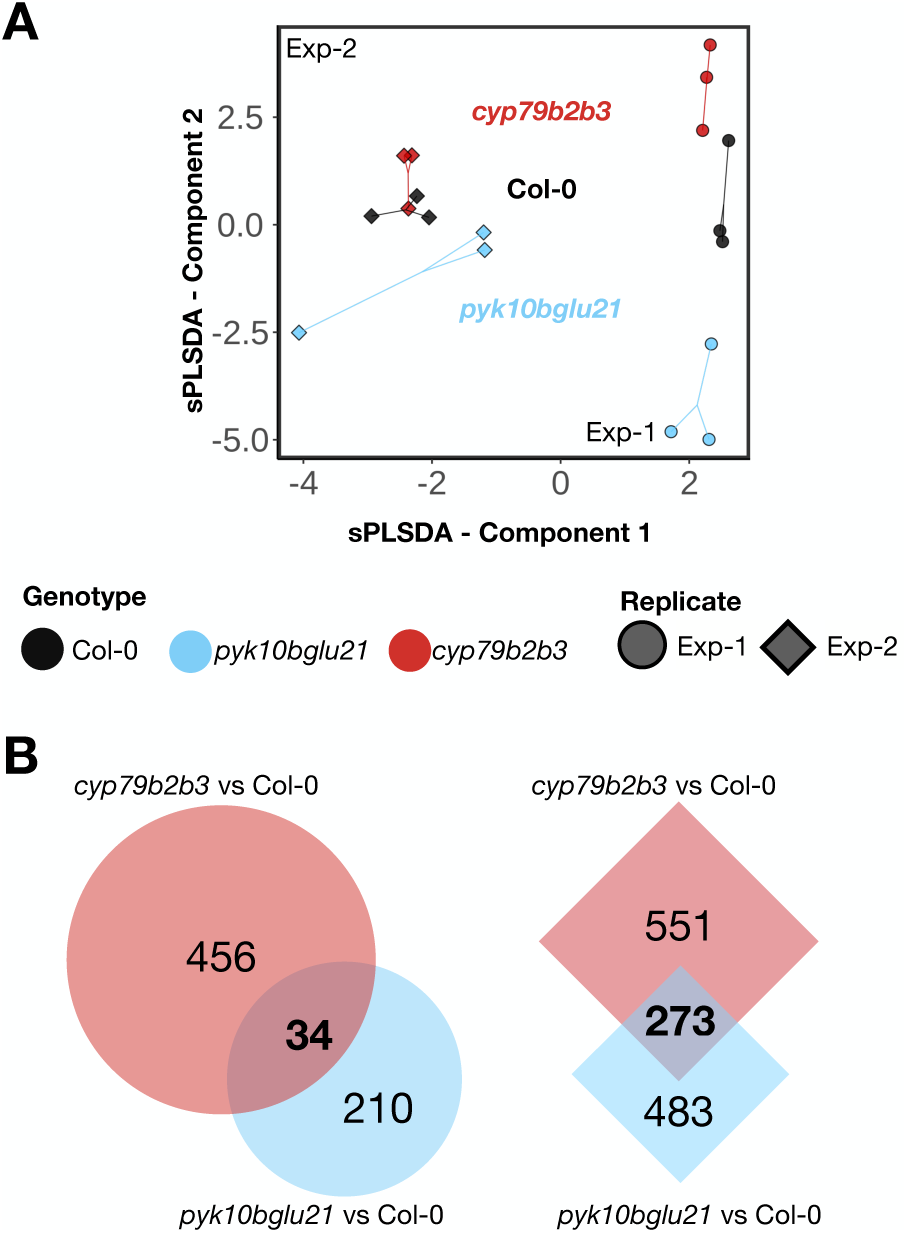
Root secretion of Trp-derived metabolites is partly dependent on PYK10 and BGLU21. (*A*) Sparse Partial Least Squares Discriminant Analysis plot of root-exuded metabolomic profiles. Colours and shapes represent the genotypes and biological replicates, respectively. (*B*) Venn diagrams representing the differentially accumulated metabolites in *cyp79b2b3* and *pyk10bglu21* mutants compared to Col-0 for each biological replicate (moderated *t* test; FDR ≤ 0.05 and |log fold change| ≥ 1.0).

Given the major variation between experimental batches, while the overall tread was similar, we decided to focus on one batch of exudate samples for further experiments. We treated natural soils with these different root exudates and analysed their impact on soil bacterial and fungal community structures (Fig. 3A). We found that, after 15 days of consecutive treatments with 2-day intervals, the soil microbial communities exhibited a shift compared to those from mock-treated soils, which is consistent with a previous study (50), in a manner dependent on the genotypes of origin of the root exudates (Fig. 3A; middle panels). Both bacterial and fungal communities in the soils treated with *pyk10bglu21* or *cyp79b2b3* exudates were significantly different from each other and from the communities in the soils treated with Col-0 exudates. Microbial taxa exhibiting differential abundance in soils treated with mutant and Col-0 root exudates were largely similar between *pyk10bglu21* and *cyp79b2b3*, (Fig S6A), and the logFC in their RA values relative to the wild-type controls were positively correlated between *pyk10bglu21* and Col-0 and *cyp79b2b3* and Col-0, both at the ASV and the family levels (Fig. 3A; right panel).

**Figure 3.**
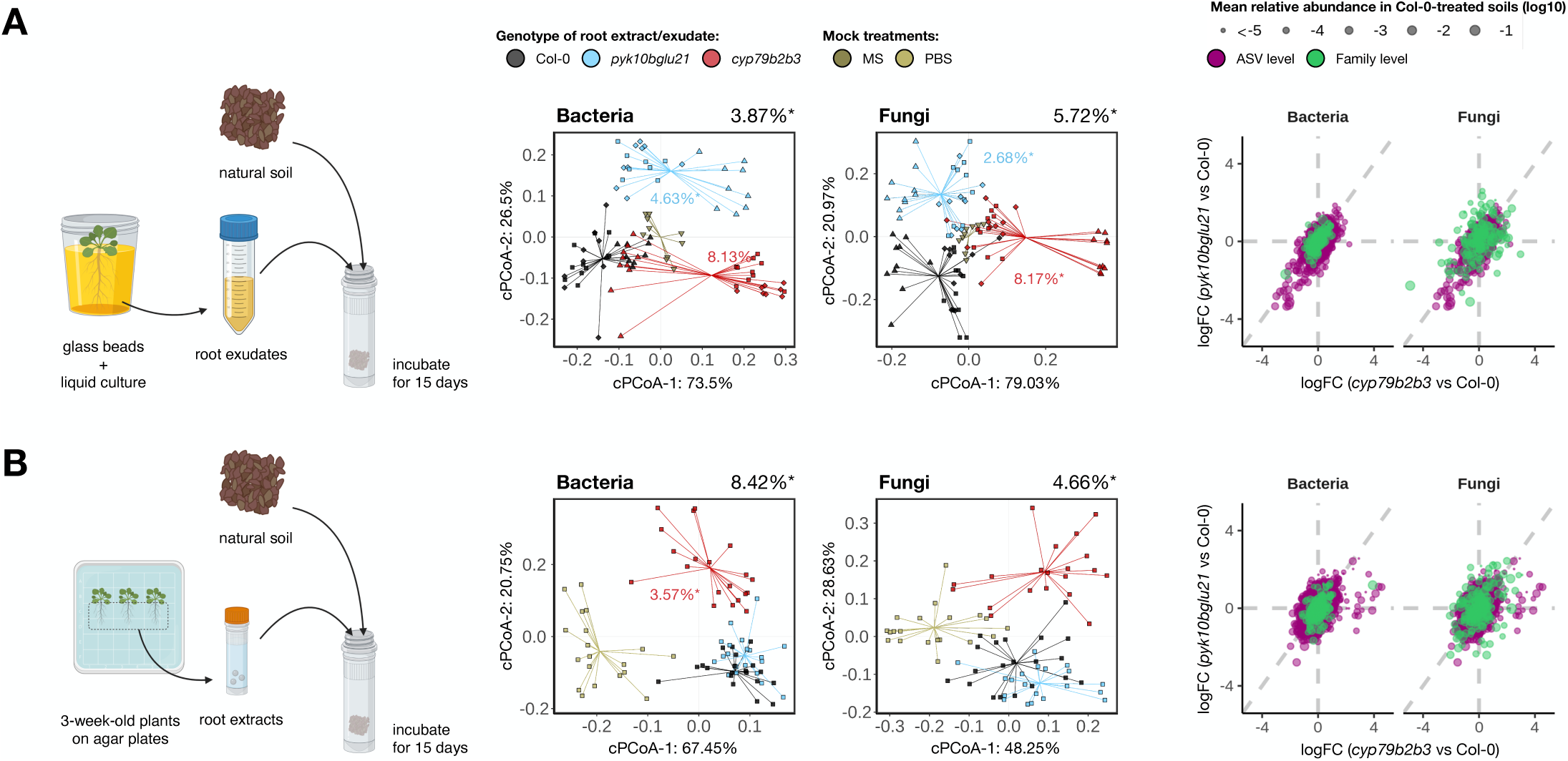
Compounds secreted into the rhizosphere in a PYK10- and BGLU21-dependent manner have an impact on microbial community assembly in the soil environment. Soils were treated with root exudates (*A*) or root extracts (*B*) consecutively for 15 days with 2-day intervals. Constrained PCoA plots are based on Bray-Curtis dissimilarities of bacterial (left) and fungal (right) communities, and the ordination was constrained by genotypes and conditioned by biological and technical replicates. The scatter plots show a comparison of changes in relative abundance of ASVs (magenta) or families (green) in soils treated with mutant root exudates/extracts compared to soils treated with Col-0 root exudates/extracts. Size corresponds to their mean relative abundance in soils treated with Col-0 root exudates/extracts.

PYK10 and BGLU21 are stored in ER bodies and do not encounter with their potential substrates in other subcellular compartments, such as vacuoles (33). Next, to test whether PYK10 and BLGLU21 contribute to the secretion of metabolites responsible for the observed community shifts or their accumulation in roots, we prepared axenic root crude extracts from these genotypes and performed the same soil treatment experiment (Fig. 3B; Table S9). The root extract treatments also triggered significant community shifts in the soil microbial community compared to mock-treated soils (Fig. 3B; middle panels). Interestingly, the constrained PCoA plots showed that Col-0 and *pyk10bglu21* root extracts triggered similar shifts in the microbial community compositions, while treatments with *cyp79b2b3* root extracts resulted in significantly different community structures. Comparison of logFC in RA values of ASVs and families also pointed to the presence of a group of microbes that were specifically affected by the CYP79B2 and CYP79B3 pathway but not by the PYK10 and BGLU21 pathway (Fig 3B, right panel; Fig S6B). These findings suggest that the Trp-derived metabolites responsible for the community assembly when treated with the Col-0 root extracts are not secreted into the rhizosphere but in fact accumulate in the *pyk10bglu21* roots. *A. thaliana* roots accumulate many myrosinases that are not stored in ER bodies, such as TGG4, TGG5, and PEN2 (51, 52), which are capable of catalysing the same reaction as PYK10 and BGLU21 when subcellular membrane partitions are artificially disrupted in a homogenate. Therefore, the active compounds may be accumulated in *pyk10bglu21* roots either in an already active form or in an inactive glycosylated form. Overall, these results suggest a role for Trp-derived metabolites whose secretion but not accumulation is dependent on ER body myrosinases in modulating microbial community structures.

### Root-exuded compounds directly affect bacterial community structure independently of fungi

Trp-derived metabolites, including IGs, have been reported to be crucial for interaction with endophytic fungi (25, 28), while their impact on bacteria remains less understood. To assess the direct impact of root exudates on bacterial community assembly, we performed a synthetic community (SynCom) profiling experiment in these root exudates (Fig. 4A), using 171 bacterial strains isolated from healthy *A. thaliana* roots grown in natural soil and 29 strains isolated from the same, unplanted soil (38). The analysis of *β*-diversity based on RA values revealed that the SynCom compositions in the Col-0 root exudates were significantly different from the composition in the mutant root exudates (Fig. 4B; Table S10). The effect on bacterial community composition was more different between Col-0 and *cyp79b2b3* exudates than between Col-0 and *pyk10bglu21* exudates, which is similar to what was observed in the soil treatment experiment (Fig. 3). The community shift in mutants compared to Col-0 was significant with both initial titres (OD_600_ of 0.05 and 0.005) and at both time points (24 and 72 hpi). The overall variance explained by genotypes was larger when we used a lower titre as a starting inoculum, and we observed a higher level of overall community growth (increase in the total number of bacterial cells within the community measured by the quantitative abundance normalized to *E. coli* cells spiked in before DNA extraction) under the low-titre than high-titre conditions, while between genotypes, the overall community growth was largely similar (Fig. S7A). When we compared the community structure dynamics over time within genotypes, we observed a more dynamic community assembly at 24 hpi when we used low-titre inocula, with eventual convergence on a similar community structure at 72 hpi (Fig. S7B). These data suggest that the contribution of ER bodies and Trp-derived specialized metabolites to community assembly is larger when bacterial cells are metabolically more active and undergoing dynamic community re-assembly. We compared the growth of each bacterial strain in Col-0 and mutant exudates (Fig. S7C) and found a small group of strains that showed a similar response in mutant exudates in comparison to Col-0 exudates (Fig. S7D), pointing to a set of bacteria commonly targeted by these pathways via root-exuded compounds. Overall, these results suggest that Trp-derived metabolites secreted to the rhizosphere in a PYK10 and BGLU21-dependent manner can directly impact the bacterial community assembly in the absence of eukaryotic organisms, such as plants and fungi.

**Figure 4.**
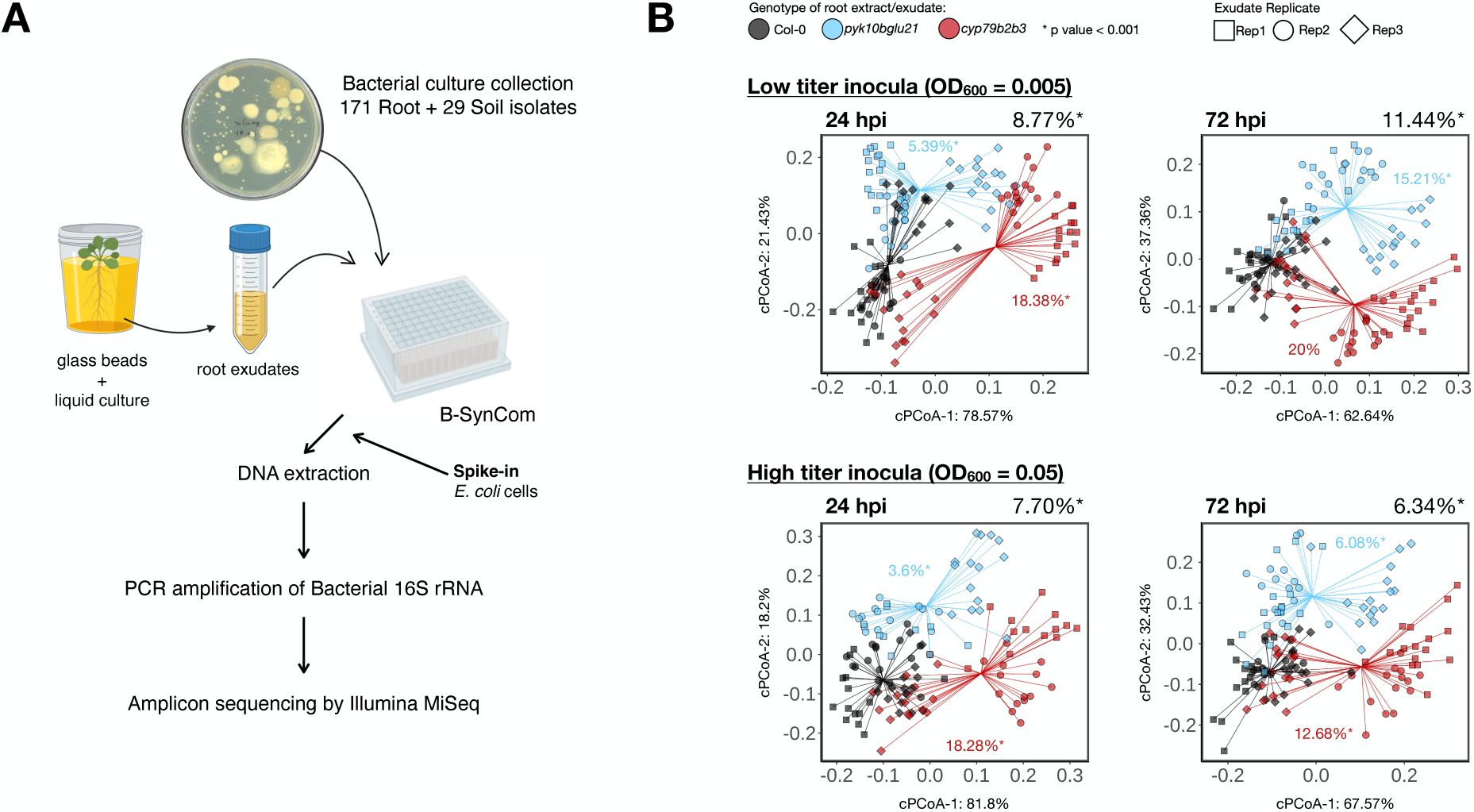
Compounds secreted into the rhizosphere in a PYK10- and BGLU21-dependent manner have a direct impact on bacterial community assembly. (*A*) Schematic representation of experiments using a bacterial SynCom composed of 171 root-derived and 29 soil-derived isolates. (*B*) Constrained PCoA analysis of bacterial communities in root exudates from Col-0 (black), *pyk10bglu21* (blue) and *cyp79b2b3* (red) at 24 and 72 hours post inoculation (hpi) using Bray-Curtis dissimilarities based on the relative abundance. Shapes correspond to the individual replicates of root exudates. Ordination was constrained by genotypes and conditioned by biological and technical replicates.

### ER bodies and Trp-derived metabolites directly influence plant-fungus interactions

Lastly, we tested whether the ER body pathway and the Trp metabolic pathway have a direct impact also on the plant-fungus interaction in the absence of bacteria. Toward this end, we inoculated the same set of plants (Col-0, *pyk10bglu21*, and *cyp79b2b3*) in a mono-association setup (24; Fig. 5) with fungal strains isolated from roots or leaves of healthy *A. thaliana* and related species grown in natural soils (25, 39). We found that the growth of *cyp79b2b3* mutant plants was severely impaired by more than half of the strains compared to the wild-type plants (24 isolates), especially by those belonging to the classes Leothiomycetes and Dothideomycetes. Furthermore, seven strains were found to significantly impair the growth of both *cyp79b2b3* and *pyk10bglu21* mutants compared to Col-0. On the other hand, 16 isolates, including all those belonging to the class Leotiomycetes, specifically restricted the growth of *cyp79b2b3* but not *pyk10bglu21* plants, and three strains showed a negative impact only on *pyk10bglu21* compared to the Col-0 wild type. These results demonstrate that the myrosinases stored in ER bodies and Trp-derived metabolites directly regulate plant-fungus interactions and target distinct but overlapping sets of fungal strains. Combined with the results that the ER body and Trp-derived metabolic pathways together contribute to the bacterial and fungal community assembly at the rhizoplane and in the endosphere, respectively, our overall findings illustrate a role for ER body-resident myrosinase-mediated Trp metabolism in root-microbiota interactions.

**Figure 5.**
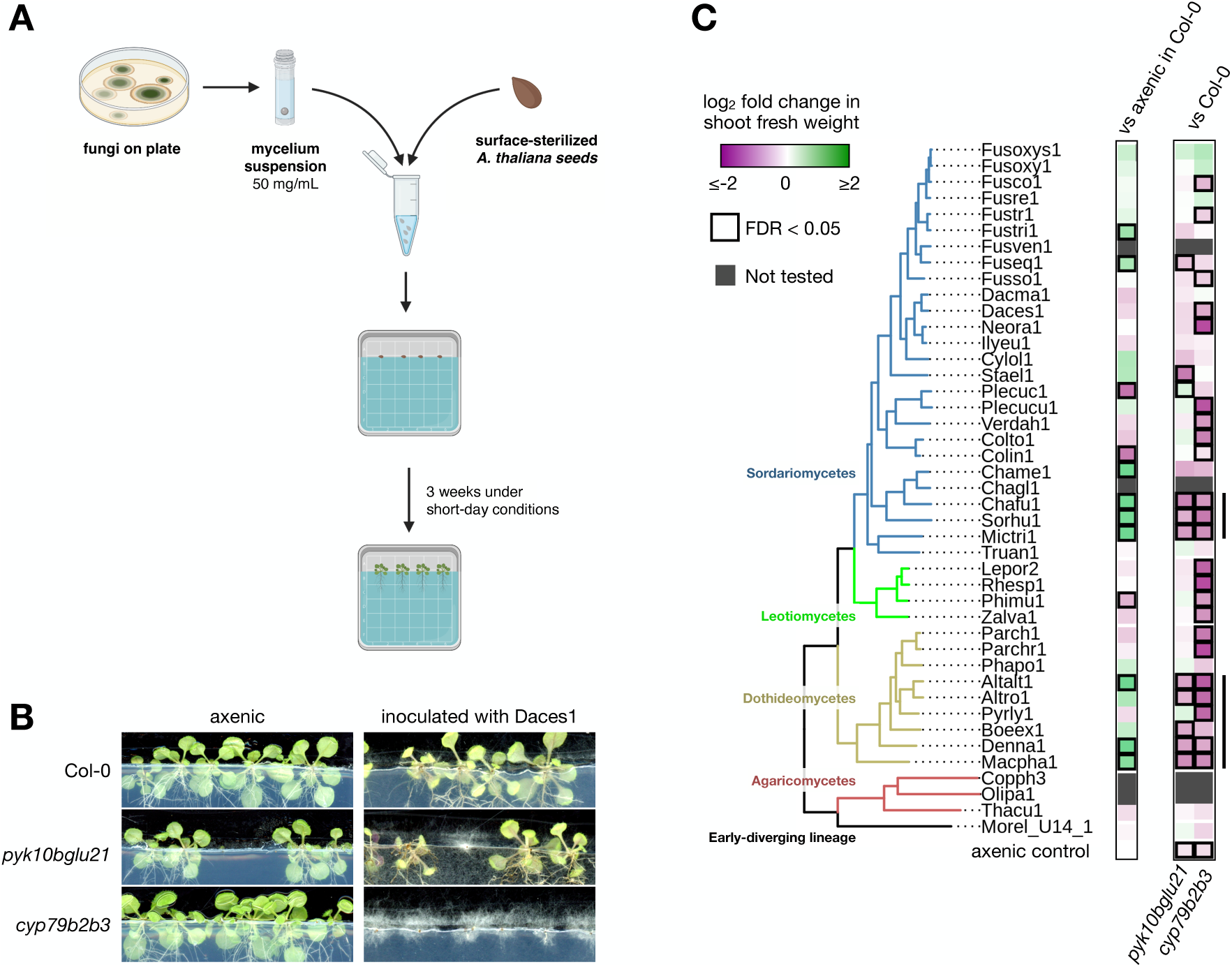
ER bodies and Trp metabolism are needed for an appropriate accommodation of endophytic fungi. (*A*) Schematic representation of experimental setup using a set of endophytic fungi isolated from healthy plant roots. (*B*) Representative images of plants inoculated with fungi along with axenic control plants. (*C*) A phylogenetic tree of fungi used in this study along with their impact on Col-0 growth compared to the axenic plants (left panel) and the impact of *pyk10bglu21* and *cyp79b2b3* mutations on shoot growth with respect to Col-0 plants inoculated with the same fungal strain. Marked with bold lines are FDR ≤ 0.05 and |logFC| ≥ 0.5.

## DISCUSSION

### The role of ER bodies and Trp-derived metabolites in root-microbiota interactions

Here, we have provided clear evidence supporting the idea that the ER body and the myrosinases that accumulate therein are involved in root-microbiota interactions, and the effects of the loss of ER body myrosinases were partly similar to the effects of the loss of Trp metabolism. This strongly suggests that there is link between ER bodies and Trp-derived compounds in interaction with soil-borne microbes. Trp metabolism produces non-glycosylated metabolites, such as camalexin and indol-3-carbonyl nitriles, and PYK10 and BGLU21 can hydrolyse glucosides derived from other amino acids, such as AGs from Met and coumarin glucosides from phenylalanine. Our pathway enrichment analysis of differentially abundant metabolites in our root exudate samples, based on the KEGG annotations of metabolites, identified biosynthetic pathways of phenylpropanoids, flavonoids and terpenoids (Table S8), without a clear commonality between *pyk10blu21* and *cyp79b2b3* exudates, further supporting the idea that the substrates of PYK10 and BGLU21 are not limited to Trp-derived glucosides. Therefore, it is plausible that the specific effects observed in each of the *pyk10bglu21* and *cyp79b2b3* mutants were due to an impaired production and/or secretion of these specific metabolites, while the common effect can be attributed to the Trp-derived glucosides, such as IGs, and their catabolites.

Of note, our metabolomic analysis of root exudates clearly indicates that PYK10 and BGLU21 contribute to the secretion of specialized metabolites, including but not limited to those derived from Trp (Fig. 2, Fig. S5, and Table S8; discussed below). This suggests that the enzymes *intracellularly* stored in ER bodies influence the composition of specialized metabolites that are *extracellularly* exuded from roots into the rhizosphere. A plausible scenario is that PYK10 and BGLU21 mediate conversion of glycosylated metabolites into a form that can be secreted into the rhizosphere, which has been reported in other plant specialized metabolites. For instance, secretion of scopoletin into the rhizosphere is severely restricted by genetic disruption of BGLU42, which is a cytosolic enzyme capable of removing the glucose moiety from scopolin, the glucosidic form of scopoletin, ultimately affecting the root-associated microbiota structure (53). We speculate that PYK10- and BGLU21-mediated removal of a glucose moiety from Trp-derived glycosylated compounds plays a role in the secretion of respective aglycones or their catabolites. Alternatively, it also remains possible that a part of PYK10 and BGLU21 proteins are secreted to the apoplast or deposited to the rhizoplane through the secretory pathway from the ER, thereby metabolizing the glucosides secreted from root tissues.

Despite the finding that the lack of PYK10 and BGLU21 showed a larger impact on the metabolomic profile of root exudates than the lack of the entire Trp metabolism (Fig. 2A), the impact on soil microbial community triggered by the lack of Trp metabolism was larger than what was triggered by the lack of these myrosinases. This suggests that, while PYK10 and BGLU21 are involved in the secretion of a wide range of metabolites under axenic conditions, a fraction of these compounds, plausibly those derived from Trp, has the predominant role in shaping the root-associated microbial community assembly.

### Possible role of IGs and IG catabolites and ER body myrosinases in root-associated microbiota assembly

Trp-derived specialized metabolites are crucial for plant-microbe interactions, and their role in plant-microbiota interactions has been well described. For example, a recent work identified a fungal dysbiosis phenotype (increased fungal load in roots) of *cyp79b2b3*, resulting in a severe plant growth defect in the presence of fungal endophytes (28). This fungal dysbiosis phenotype was not observed in the *myb34/51/122* mutant roots or in any known mutants in the Trp-derived branch pathways, pointing to the presence of uncharacterized metabolites that are crucial for regulating the fungal load in roots. In this study, we observed a significant alteration in the bacterial and fungal communities in roots of *myb34/51/122*, which is specifically impaired in IG biosynthesis but retains the ability to produce other Trp-derived metabolites (35), indicating that IGs also play a role in root-microbiota interactions. On the other hand, we did not detect substantial amounts of intact IGs in the root exudates, even those collected from Col-0 wild-type plants (Figure S5L-N), which is consistent with what was reported in a previous study employing a similar experimental approach (30). Low levels of IGs in root exudates could be due to rapid degradation of IGs after root secretion by an uncharacterized extracellular myrosinase or a requirement for glucose removal for secretion of IG catabolites. Alternatively, it is also possible that IGs and their catabolites are secreted at lower levels than expected from axenic and naïve plants under our experimental conditions without any immuno-stimuli, as an activation of immune responses by microbe-associated molecular patterns (MAMPs) induces secretion of Trp-derived metabolites including camalexin (26). Because several IG catabolites, such as isothiocyanates and carbinols, are unstable and difficult to detect, and our understanding of other stable IG catabolites remains limited, it remains to be determined whether IGs and IG catabolites are secreted but not detected or whether these compounds are not secreted in substantial amounts into the rhizosphere.

### Auxin and other Trp-derived metabolites can be secreted in an ER body myrosinase-dependent manner and contribute to bacterial community assembly

Our data also do not exclude the possibility that PYK10 and BGLU21 influence secretion of Trp-derived metabolites other than IGs that play a key role in manipulating microbial community composition. We detected a variety of Trp-derived specialized metabolites in our exudate samples, such as indole-3-acetic acid (IAA), indole-3-acetonitrile (IAN) and indole-3-carboxylic acid (ICA) (Fig. S5). Of note, the RA values of IAA in exudates collected from *pyk10bglu21* and *cyp79b2b3* mutant roots were lower than in exudates from Col-0 roots (*P* < 0.001 for *cyp79b2b3* vs Col-0; *P* = 0.0516 for *pyk10bglu21*), and a similar trend was observed in IAN (Fig. S5). Trp metabolism is initiated with oxidation of Trp by CYP79B2 and CYP79B3, producing IAOx as a product. IAOx is further oxidized to IAN or 1-aci-nitro-2-indolyl-ethane by CYP71A12 and CYP71A13 or CYP83B1, respectively, which are the precursors of camalexin and IGs, respectively. IAN can also be a substrate for a group of nitrilases that directly produce IAA (54), which is one of the multiple Trp-derived IAA biosynthetic pathways (37). Importantly, IAN can also be produced as a terminal product of IG catabolic processes mediated by myrosinases, and a recent simulation modelling study suggested that plants can alter IAA signalling dynamics following the hydrolysis of IGs in a manner dependent on the responsible IG-hydrolysing myrosinases (55). Therefore, it is plausible that the hydrolysis of IGs by PYK10 and BGLU21 contributes to the secretion of IAA into the rhizoplane/rhizosphere. Auxin, including IAA, by itself has an impact on microbial behaviour (56, 57), and root-associated commensal bacteria are capable of modulating auxin accumulation in roots (58). Overall, these data raise the possibility that modulation of the microbial community by root exudates is partly accounted for by auxin and that the root secretion of auxin is directly or indirectly dependent on the myrosinase activity of PYK10 and BGLU21.

It is also important to note that the exudates used in our experiments were collected from axenic, naïve plants, i.e., in the absence of microbes or any immune elicitors. It has been well described that plant immune status has an impact on root metabolomic profiles (26, 52) as well as on root exudation profiles (59–61). It is therefore likely that the metabolomic profile of root exudates may be substantially different between axenic and soil conditions. Our results nonetheless provide evidence that Trp-derived metabolites that are secreted in a manner dependent on PYK10 and BGLU21 under axenic naïve conditions, including IAA and IAN, are capable of manipulating a microbial community; further studies will address to which extent our insights into the activity of axenic root exudates can explain the actual microbial community shift observed in the greenhouse experiment (Fig. 1).

### Root-produced glucosides might represent a general plant strategy to shape the soil-borne microbiome

Glucosinolates and ER bodies are specifically found in plants within the order Brassicales and represent lineage-specific innovations of plant secondary metabolism and its regulation, respectively. While IGs are conserved across the entire order (62), the ER body system appears to be a more recent innovation, arising approximately between 50 and 60 million years ago (63), just before separation of Brassicacaee, Capparaceae, and Cleomaceae from the other Brassicales (64). This suggests that ER bodies contribute to boosting the overall efficacy of the IG-mediated defence system. Such lineage-specific innovation of glucosides that are important for plant-microbe interactions appears to be a common feature of plants. For instance, benzoxazinoids in maize are crucial for rhizosphere microbiota assembly (65), and benzoxazinoids are also produced and accumulate as glucosides that are activated by the action of respective BGLUs (66). The role of benzoxazinoids in plant-soil feedback indicates the involvement of root exudates; whether it is benzoxazinoids or their glycosylated forms that are secreted into soil remains unclear. Glycosylation of bioactive specialized metabolites to avoid autotoxicity appears to be a common strategy for plants, which enables rapid responses to environmental cues, such as pathogenic/herbivorous invasion (67). Of note, it is proposed that glucosinolates arose from cyanogenic glycosides, which are also important for anti-microbial defence (68) and undergo very similar metabolic processing during their activation, including hydrolytic deglycosylation by cyanogenic BGLUs and detoxification by glutathione conjugation (69). Given the toxicity of HCN that is formed during the deglycosylation of cynogenic glucosides, it is possible that cyanogenic glycosides are also involved in root microbiota assembly. Together with the fact that BGLU activity is needed for secretion of scopoletin into the rhizosphere via removal of glucose from its glycosylated from, scopolin (70), glucose conjugation may be important not only for suppression of toxicity but also to control secretion processes. Based on these insights, we propose that root microbiota assembly mediated by root-produced glucosides and the facilitation of their bioactivity and/or their secretion by cognate BGLUs is a widespread strategy in plants to modulate root microbiota compositions, albeit exploiting different classes of glucosides depending on the plant lineages.

### Data Availability

The raw sequences are available at the European Nucleotide Archive (ENA) under the accession number PRJEB54088. The scripts used for processing the Illumina reads and statistical analyses are available at https://github.com/Guan06/DADA2_pipeline and https://github.com/arpankbasak/ERBody_RootMicrobiota.

## Supplementary data

**Figure S1.** Community shifts observed in the mutant roots compared to Col-0 roots retained at the family level.

**Figure S2.** Similar effects of ER body pathway and Trp metabolism on root microbiota community structure at the ASV level.

**Figure S3.** Similar effects of ER body pathway and Trp metabolism on root microbiota community structure at the family level.

**Figure S4.** Several bacterial and fungal families are commonly enriched or depleted in mutant roots compared to wild-type roots.

**Figure S5.** Composition of specialized metabolites differing within the mutant root exudates.

**Figure S6.** The relative abundance of microbes is different in the mutant root compartment compared to the wild type.

**Figure S7.** Community dynamics and growth of individual microbes in a SynCom treated with root exudates.

**Table S1.** List of bacterial strains used in the SynCom.

**Table S2.** List of fungal strains used in the mono-association assay.

**Table S3.** Barcoded primers used for bacterial 16s V5–V7 amplicon sequencing.

**Table S4.** Barcoded primers used for bacterial ITS1 amplicon sequencing.

**Table S5.** Forward barcoded primer set for bacterial 16s V5–V7 amplicon sequencing.

**Table S6.** Forward barcoded primer set for bacterial ITS1 amplicon sequencing.

**Table S7.** Summary statistics of community structure analysis by constrained ordination followed by pairwise PERMANOVA for the greenhouse experiment.

**Table S8.** Summary pathway enrichment analysis of root exudate metabolome.

**Table S9.** Summary statistics of community structure analysis by constrained ordination followed by pairwise PERMANOVA for the soil treatment experiment.

**Table S10.** Summary statistics of community structure analysis by constrained ordination followed by pairwise PERMANOVA for the SynCom experiment.

## Acknowledgements

We thank Paul Schulze-Lefert for providing all necessary infrastructures and instrumentation at the Max Planck Institute for Plant Breeding Research, as well as Małopolska Centre of Biotechnology, Jagiellonian University, for their institutional support. We are grateful to Anna Lisa Roth, Brigitte Pickel, Dieter Becker and Diana Dresbach for their technical support and to Kathrin Wippel, Paloma Durán, and Tomohisa Shimasaki for their technical help in the hydroponic culture, community profiling and soil treatment experiments, respectively. We thank Neysan Donnelly for their help in editing the manuscript.

## Funding

This work was supported by an Overseas Research Fellowship funded by the Japanese Society for the Promotion of Science (JSPS) to R.T.N., the Priority Programme “Reconstruction and Deconstruction of Plant Microbiota” (DECRyPT) [402201269] funded by the Deutsche Forschungsgemeinschaft (DFG) to R.T.N., the National Science Centre of Poland grants OPUS [UMO-2016/23/B/NZ1/01847] to A.K.B. and K.Y. and HARMONIA [UMO-2015/18/M/NZ1/00406] to P.B.

## Supplementary figure legends

**Figure S1. Community shifts observed in the mutant roots compared to Col-0 roots retained at the family level.** Constrained principal coordinates analysis (PCoA) of the bacterial and fungal community structures in the roots of Col-0 as well as mutants impaired in ER body formation (*nai1-1*), ER body-accumulating myrosinases (*pyk19bglu21*), IG biosynthesis (*myb34/51/122*), and Trp metabolism (*cyp79b2b3*) based on Bray-Curtis dissimilarities computed from the relative abundance aggregated at the family level. Ordination was constrained by genotypes and conditions by soil batches, biological replicates and sequencing runs. Colours and shapes represent the genotypes and biological replicates, respectively. Variation explained by genotypes and respective *P* values based on permutational analyses of variance (PERMANOVA; n = 999) are indicated at top-left.

**Figure S2. Similar effects of ER body pathway and Trp metabolism on root microbiota community structure at the ASV level.** Comparison of log_2_-scale fold changes in relative abundance of bacterial ASVs (A and B) and fungal ASVs (C and D) in rhizoplane (A and C) and endosphere (B and D) fractions of mutants compared to respective Col-0. ASVs that are consistently detected in all genotypes are marked with solid lines. Open and closed points correspond to two independent soil batches. Pearson’s correlation coefficients are indicated at top-left.

**Figure S3. Similar effects of ER body pathway and Trp metabolism on root microbiota community structure at the family level.** Comparison of log_2_-scale fold changes in relative abundance of bacterial ASVs (A and B) and fungal ASVs (C and D) aggregated at the family level in rhizoplane (A and C) and endosphere (B and D) fractions of mutants compared to respective Col-0. ASVs that are consistently detected in all genotypes are marked with solid lines. Open and closed points correspond to two independent soil batches. Pearson’s correlation coefficients are indicated at top-left.

**Figure S4. Several bacterial and fungal families are commonly enriched or depleted in mutant roots compared to wild-type roots**. The dotted heatmap represents log_2_-scale fold changes in relative abundance of bacterial (A) and fungal ASVs (B) aggregated at the family level in mutant roots compared to Col-0 roots. The mean aggregated relative abundance of each family across all genotypes in each soil batch is shown as a barplot.

**Figure S5. Relative amounts of glucosinolates, auxin, and coumarins in the mutant root exudates.** Relative abundance of metabolites in exudates normalized to Col-0 root exudates are shown as boxplots. Aliphatic (A–J), benzyl (K) and indole glucosinolates (L–N), as well as known indolic compounds (O and P) and coumarins (Q–S) are quantified based on either standards or KEGG annotation. Letters indicate statistical significance corresponding to ANOVA and post-hoc Tukey’s HSD tests within each metabolite (*α* = 0.05). Metabolites without statistical significance based on ANOVA are shown without letters. 9MSN, 9- methylsulfinylnonyl glucosinolate; 9MTN, 9-methylthiononyl glucosinolate; 5MTP, 5- methylthiopentyl glucosinolate; 8MSO, 8-methylsulfinyloctly glucosinolate; 3MSB, 3- methylsulfinylpropyl glucosinolate; 6MTH, 6-methylthiohexyl glucosinolate; 10MSD, 10- methylsulfonyldecyl glucosinolate; 1MI3G, 1-methoxyindol-3-ylmethyl glucosinolate; 4MI3G, 4- methoxyindol-3-ylmethyl glucosinolate; I3G, indol-3-ylmethyl glucosinolate; IAN, indole-3- acetonitrile; IAA, indole-3-acetic acid.

**Figure S6. The relative abundance of microbes is different in the mutant root compartment compared to the wild type**. The dotted heatmap represents log_2_-scale fold changes in relative abundance of bacterial (A) and fungal ASVs (B) aggregated at the family level in soils treated with mutant root exudates or extracts compared to the soils treated with Col-0 root exudates or extracts. The mean aggregated relative abundance of each family across all genotypes in each treatment is shown as a barplot.

**Figure S7. Community dynamics and growth of individual microbes in a SynCom treated with root exudates.** (A) The overall growth of the 200-member SynCom based on the aggregated quantitative abundance (QA) of the strains relative to spiked-in DH5*α*. Top panels, experiment 1; bottom panels, experiment 2. Left panels, low-titre inocula (OD_600_ = 0.005); right panels, high-titre inocula (OD_600_ = 0.05). Colours represent the genotypes from which root exudates were collected. (B) Constrained PCoA analysis of SynCom that compared community structures when treated with the root exudates from the same genotype, based on Bray-Curtis dissimilarities computed from relative abundances. Ordinations are constrained by the starting titres and time points (represented by colours) and conditioned by technical and biological replicates as well as independent batches of root exudate (represented by shapes). Arrows indicate the community shift over the incubation period. Numbers on top indicate the overall variance explained by the initial titre and the time points. (C) Heatmaps showing the taxonomy, quantitative abundance relative to spiked-in DH5*α* and log2-scale fold change of QA (QA logFC) in mutant root exudates compared to Col-0 root exudates. (D) Comparison of QA logFC in mutant exudates compared to Col-0 exudates of the strains whose mean QA is higher than 5. Strains whose growth is commonly promoted or suppressed in both mutant exudates are represented by green or magenta, respectively.

## Notes

The authors declare no conflict of interest.

### Competing Interest Statement

The authors have declared no competing interest.

