## Supplementary Figures for "Tryptophan specialized metabolism and ER body-resident myrosinases modulate root microbiota assembly"

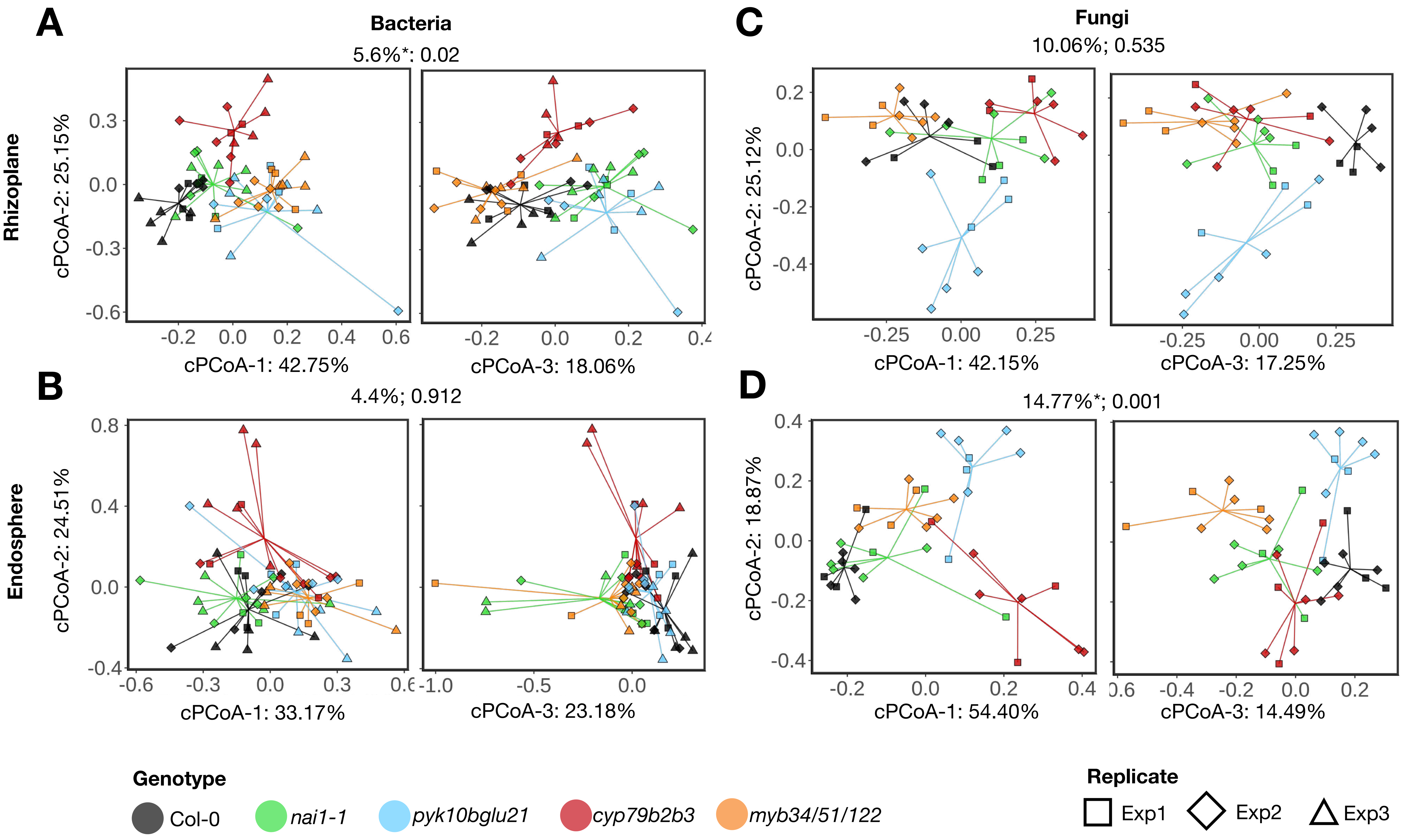


**Figure S1. Community shifts observed in the mutant roots compared to Col-0 roots retained at the family level.** Constrained principal coordinates analysis (PCoA) of the bacterial and fungal community structures in the roots of Col-0 as well as mutants impaired in ER body formation (*nai1-1*), ER body-accumulating myrosinases (*pyk19bglu21*), IG biosynthesis (*myb34/51/122*), and Trp metabolism (*cyp79b2b3*) based on Bray-Curtis dissimilarities computed from the relative abundance aggregated at the family level. Ordination was constrained by genotypes and conditions by soil batches, biological replicates and sequencing runs. Colours and shapes represent the genotypes and biological replicates, respectively. Variation explained by genotypes and respective *P* values based on permutational analyses of variance (PERMANOVA; n = 999) are indicated at top-left.

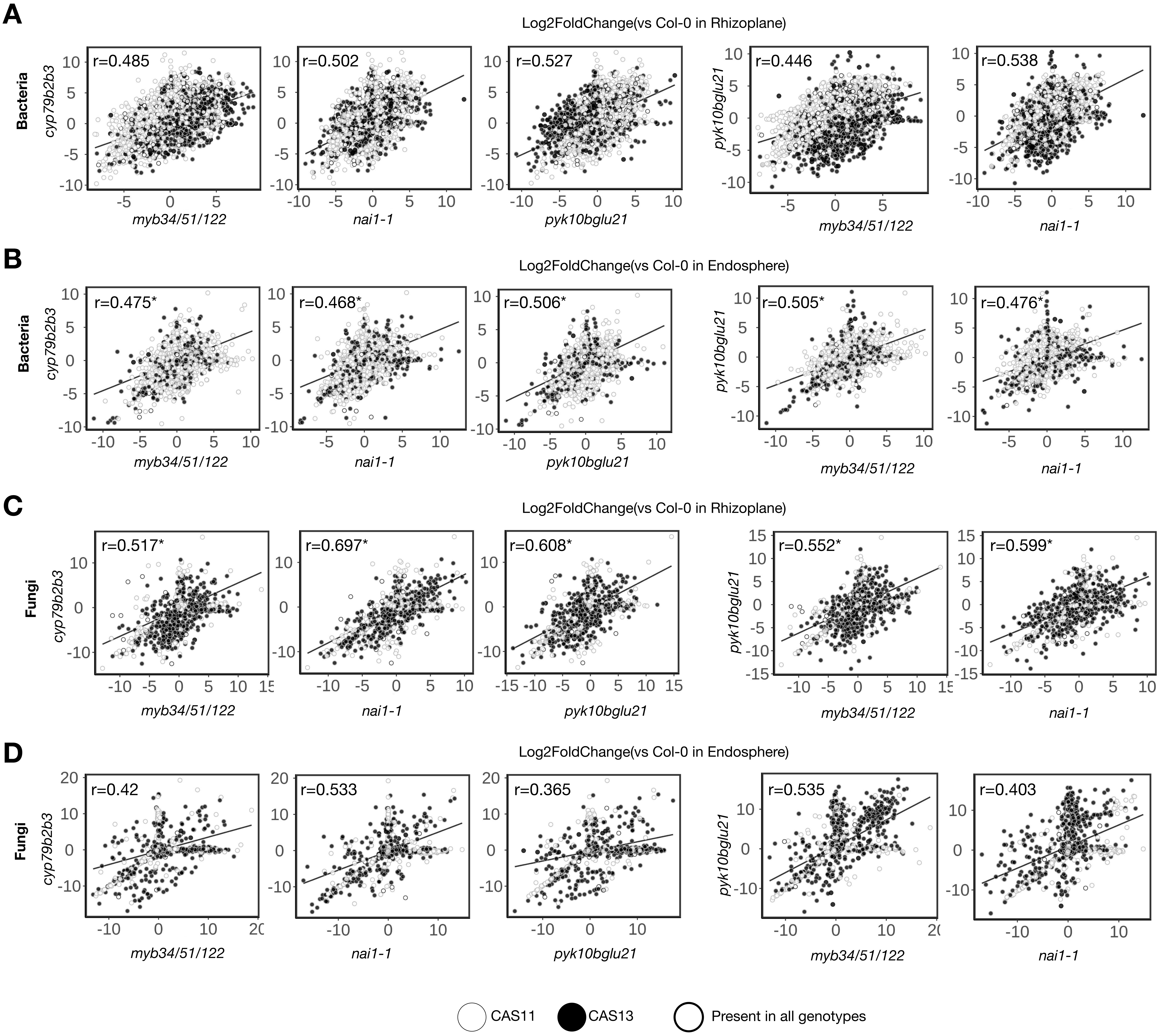


**Figure S2. Similar effects of ER body pathway and Trp metabolism on root microbiota community structure at the ASV level.** Comparison of log$_2$-scale fold changes in relative abundance of bacterial ASVs (A and B) and fungal ASVs (C and D) in rhizoplane (A and C) and endosphere (B and D) fractions of mutants compared to respective Col-0. ASVs that are consistently detected in all genotypes are marked with solid lines. Open and closed points correspond to two independent soil batches. Pearson's correlation coefficients are indicated at top-left.

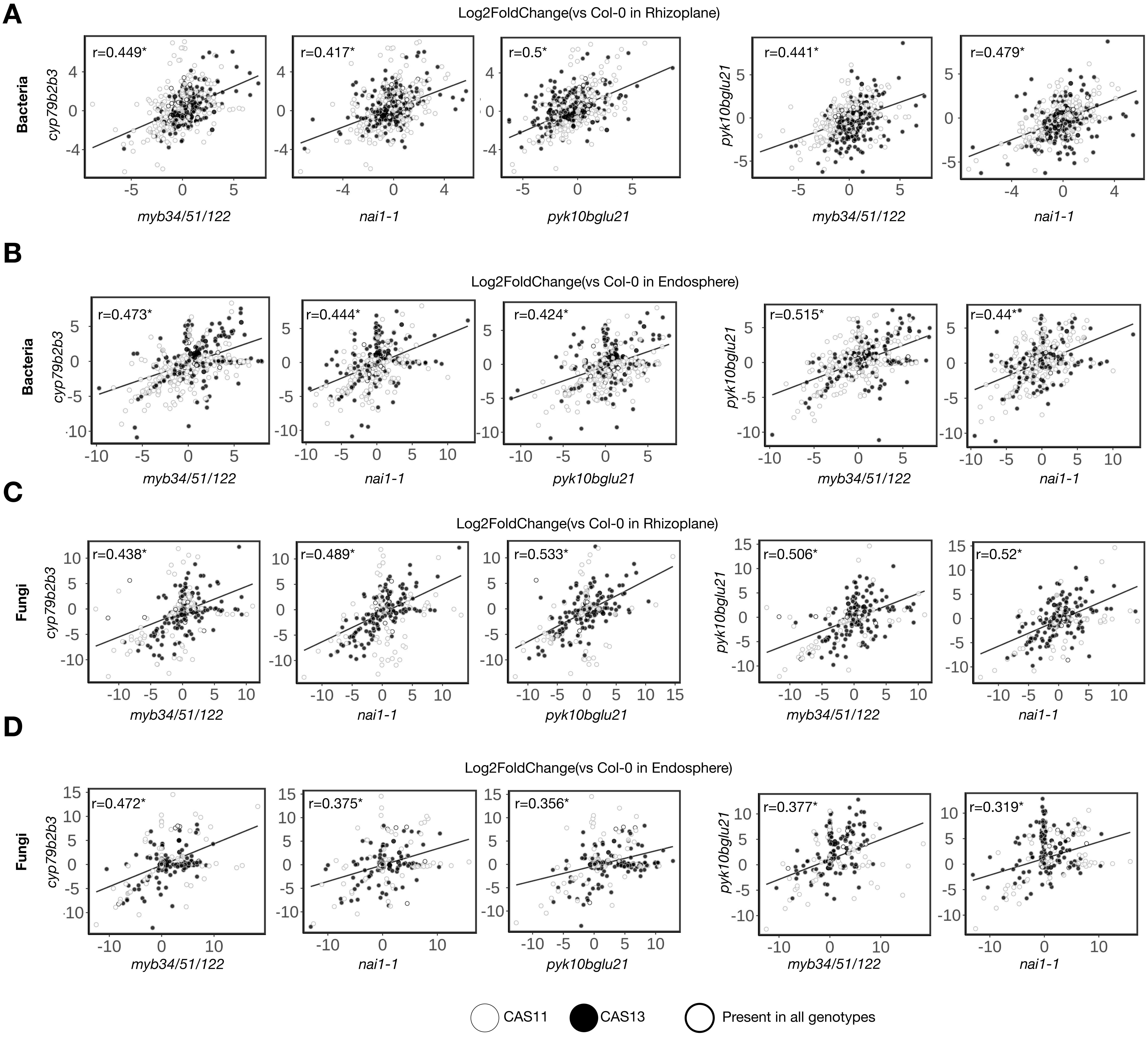


**Figure S3. Similar effects of ER body pathway and Trp metabolism on root microbiota community structure at the family level.** Comparison of $log_2$-scale fold changes in relative abundance of bacterial ASVs (A and B) and fungal ASVs (C and D) aggregated at the family level in rhizoplane (A and C) and endosphere (B and D) fractions of mutants compared to respective Col-0. ASVs that are consistently detected in all genotypes are marked with solid lines. Open and closed points correspond to two independent soil batches. Pearson's correlation coefficients are indicated at top-left.

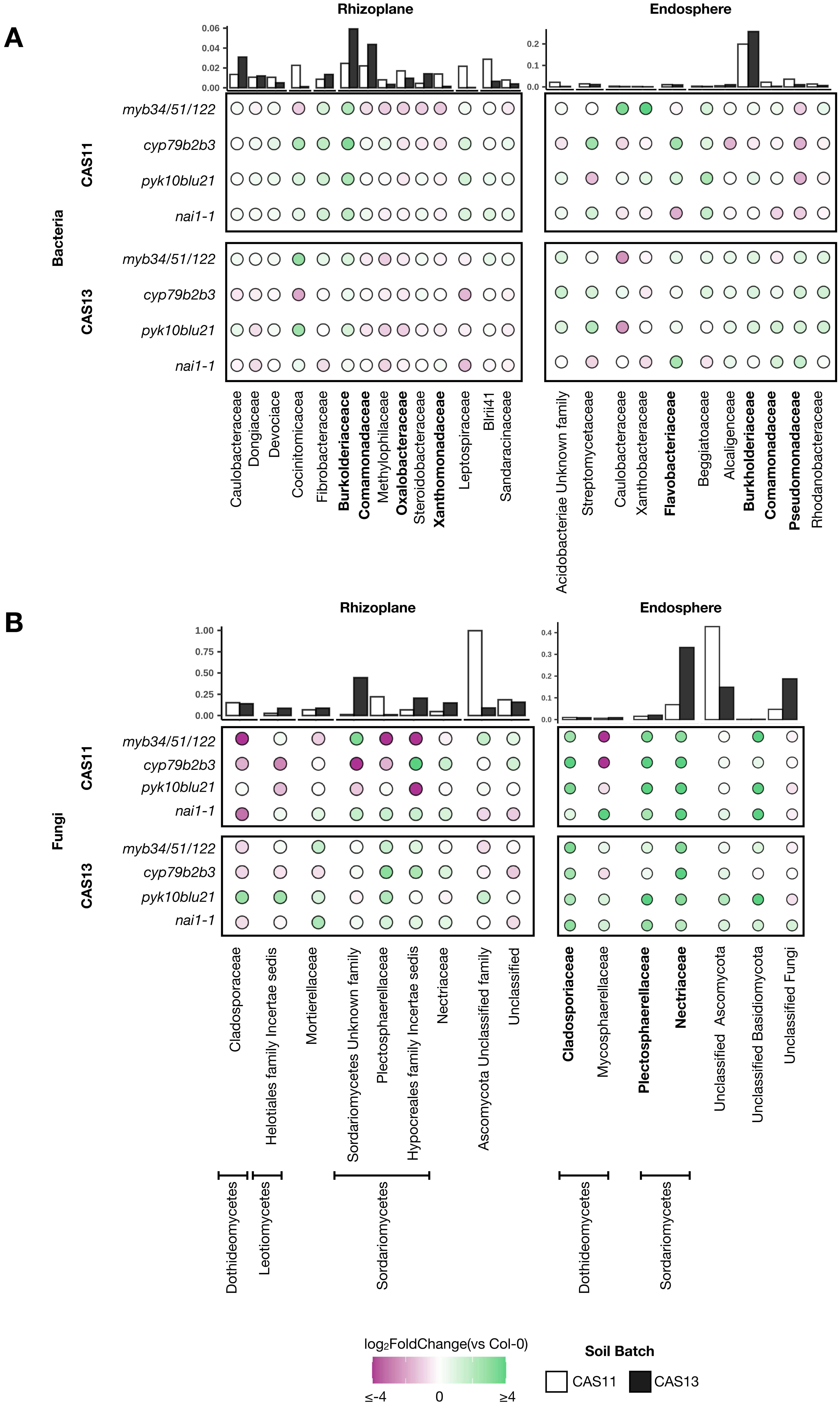


**Figure S4. Several bacterial and fungal families are commonly enriched or depleted in mutant roots compared to wild-type roots**. The dotted heatmap represents $\log_2$-scale fold changes in relative abundance of bacterial (A) and fungal ASVs (B) aggregated at the family level in mutant roots compared to Col-0 roots. The mean aggregated relative abundance of each family across all genotypes in each soil batch is shown as a barplot.

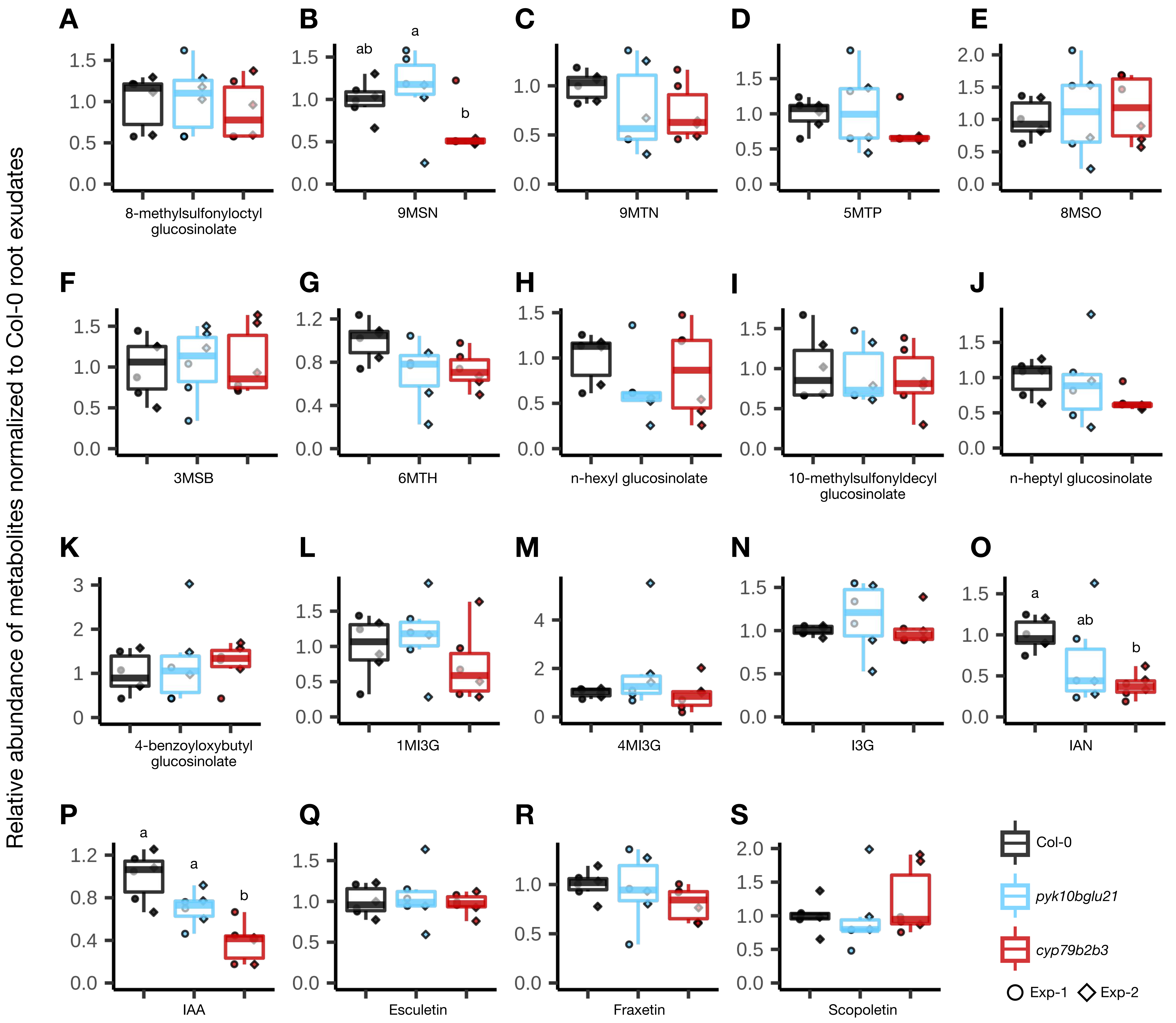


**Figure S5. Relative amounts of glucosinolates, auxin, and coumarins in the mutant root exudates.** Relative abundance of metabolites in exudates normalized to Col-0 root exudates are shown as boxplots. Aliphatic (A–J), benzyl (K) and indole glucosinolates (L–N), as well as known indolic compounds (O and P) and coumarins (Q–S) are quantified based on either standards or KEGG annotation. Letters indicate statistical significance corresponding to ANOVA and post-hoc Tukey's HSD tests within each metabolite (a = 0.05). Metabolites without statistical significance based on ANOVA are shown without letters. 9MSN, 9-methylsulfinylnonyl glucosinolate; 9MTN, 9-methylthiononyl glucosinolate; 5MTP, 5-methylthiopentyl glucosinolate; 8MSO, 8-methylsulfinyloctly glucosinolate; 3MSB, 3-methylsulfinylpropyl glucosinolate; 6MTH, 6-methylthiohexyl glucosinolate; 10MSD, 10-methylsulfonyldecyl glucosinolate; 1MI3G, 1-methoxyindol-3-ylmethyl glucosinolate; 4MI3G, 4-methoxyindol-3-ylmethyl glucosinolate; I3G, indol-3-ylmethyl glucosinolate; IAN, indole-3-acetonitrile; IAA, indole-3-acetic acid.

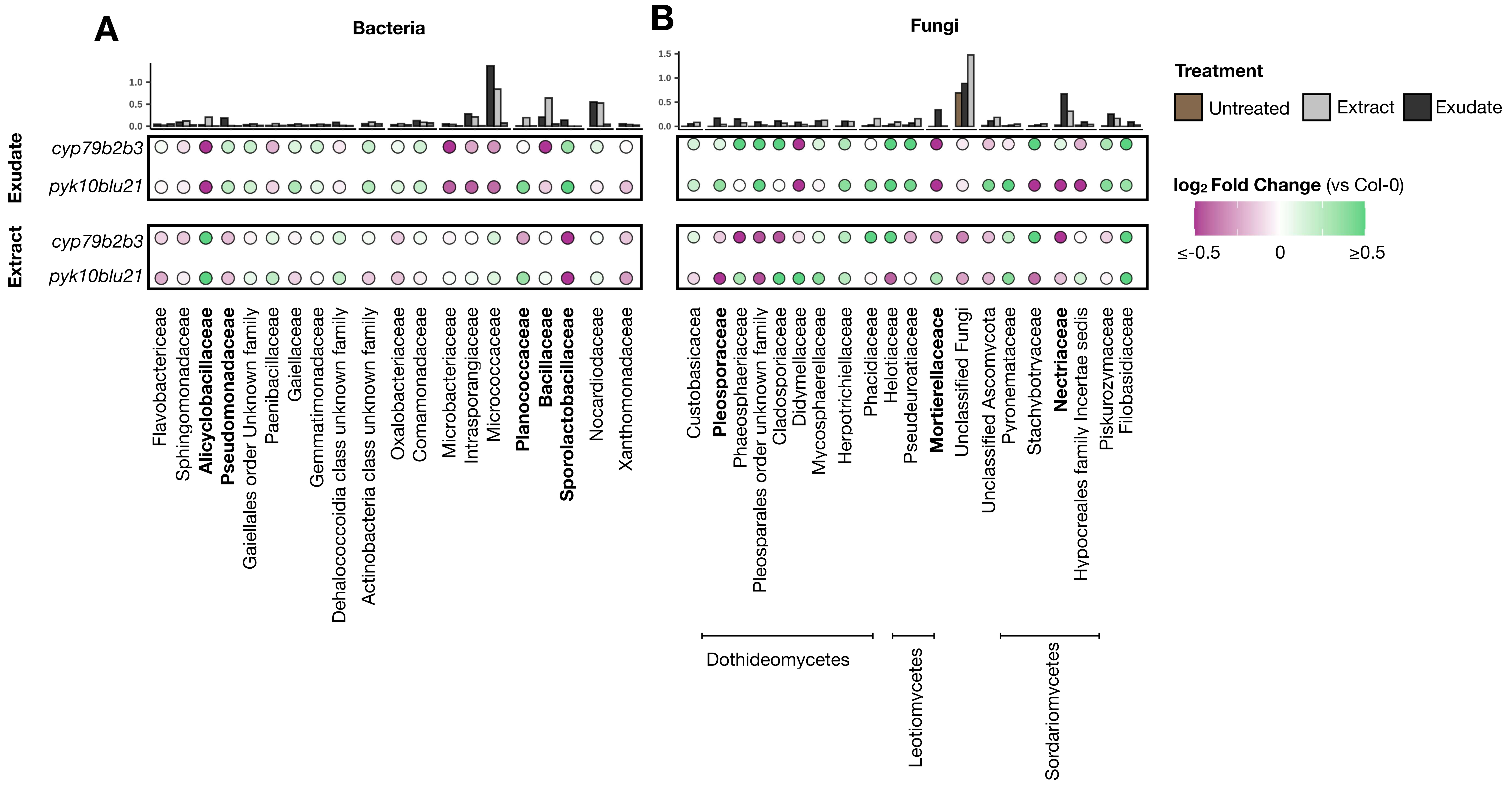


**Figure S6. The relative abundance of microbes is different in the mutant root compartment compared to the wild type**. The dotted heatmap represents $log_2$-scale fold changes in relative abundance of bacterial (A) and fungal ASVs (B) aggregated at the family level in soils treated with mutant root exudates or extracts compared to the soils treated with Col-0 root exudates or extracts. The mean aggregated relative abundance of each family across all genotypes in each treatment is shown as a barplot.

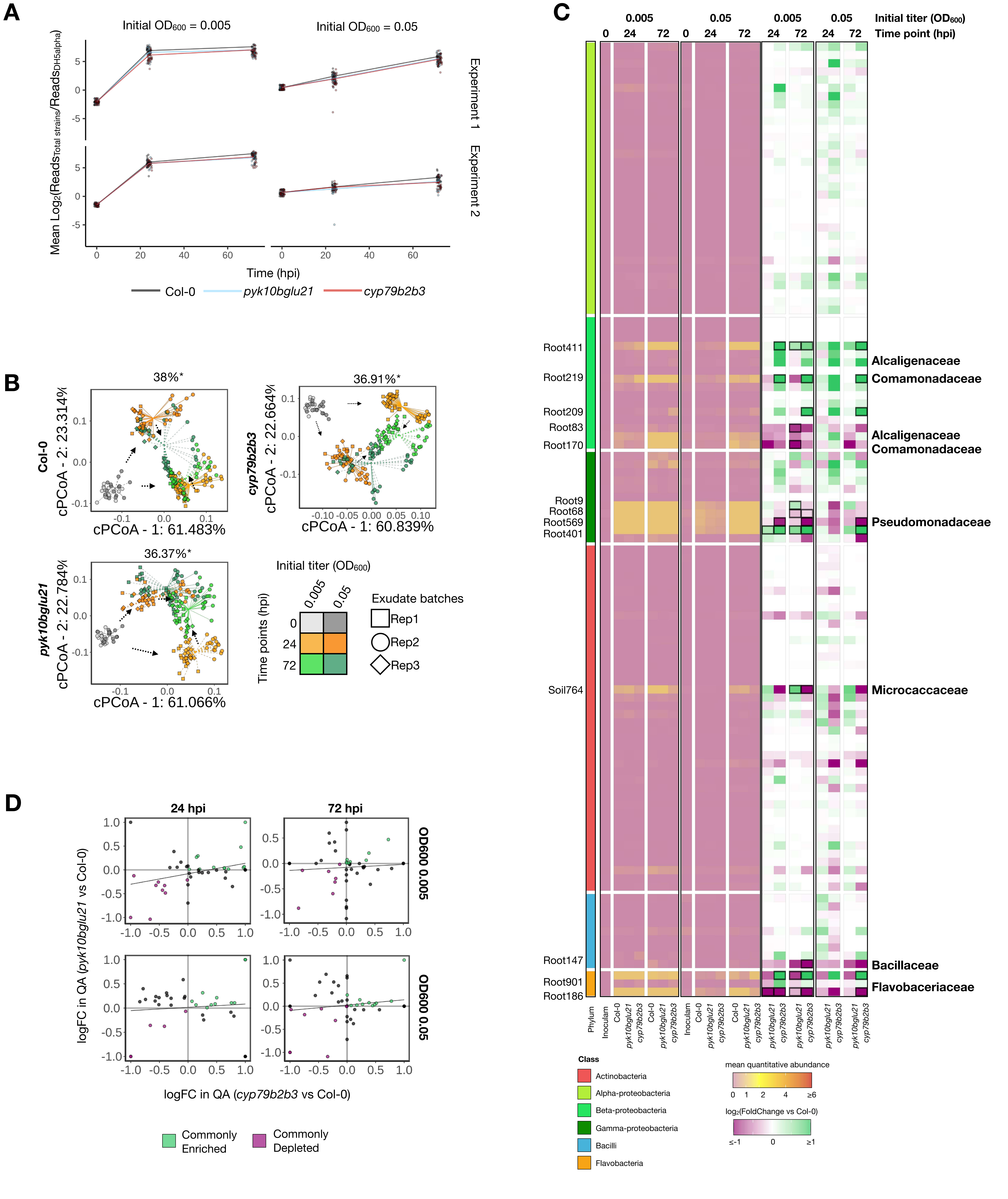


**Figure S7. Community dynamics and growth of individual microbes in a SynCom treated with root exudates.** (A) The overall growth of the 200-member SynCom based on the aggregated quantitative abundance (QA) of the strains relative to spiked-in DH5a. Top panels, experiment 1; bottom panels, experiment 2. Left panels, low-titre inocula ($OD_{600}$ = 0.005); right panels, high-titre inocula ($OD_{600}$ = 0.05). Colours represent the genotypes from which root exudates were collected. (B) Constrained PCoA analysis of SynCom that compared community structures when treated with the root exudates from the same genotype, based on Bray-Curtis dissimilarities computed from relative abundances. Ordinations are constrained by the starting titres and time points (represented by colours) and conditioned by technical and biological replicates as well as independent batches of root exudate (represented by shapes). Arrows indicate the community shift over the incubation period. Numbers on top indicate the overall variance explained by the initial titre and the time points. (C) Heatmaps showing the taxonomy, quantitative abundance relative to spiked-in DH5a and log2-scale fold change of QA (QA logFC) in mutant root exudates compared to Col-0 root exudates. (D) Comparison of QA logFC in mutant exudates compared to Col-0 exudates of the strains whose mean QA is higher than 5. Strains whose growth is commonly promoted or suppressed in both mutant exudates are represented by green or magenta, respectively.
